# Metabolomics reveals that dietary xenoestrogens alter cellular metabolism induced by palbociclib/letrozole combination cancer therapy

**DOI:** 10.1101/188102

**Authors:** Benedikt Warth, Philipp Raffeiner, Ana Granados, Tao Huan, Mingliang Fang, Erica M Forsberg, H. Paul Benton, Laura Goetz, Caroline H. Johnson, Gary Siuzdak

## Abstract

**Highlights:**

- Synergism of combined palbociclib/letrozole chemotherapy was examined by global metabolomics
- Combination therapy led to more pronounced effects on the MCF-7 metabolome than single agents
- Dietary phyto- and mycoestrogens significantly affected the metabolic and anti-oncogenic response of the drugs
- Implications of these bio-active chemicals on therapeutic success in breast cancer patients appear plausible

**In Brief:** Warth et al. used innovative global metabolomics and pathway prediction technology to describe the metabolic effects of the combined palbociclib/letrozole breast cancer therapy. Moreover, the role of dietary xenoestrogens on this treatment was examined by metabolite data, proliferation experiments, and functional assays.

**Summary:** Recently, the palbociclib/letrozole combination therapy was granted accelerated FDA approval for the treatment of estrogen receptor (ER) positive breast cancer. Since the underlying metabolic effects of these drugs are yet unknown, we investigated their synergism at the metabolome level in MCF-7 cells. As xenoestrogens interact with the ER, we additionally aimed at deciphering the impact of the phytoestrogen genistein, and the estrogenic mycotoxin zearalenone on this treatment. A global metabolomics approach was applied to unravel metabolite and pathway modifications. The results clearly showed that the combined effects of palbociclib and letrozole on cellular metabolism were far more pronounced than that of each agent alone and potently influenced by xenoestrogens. This behavior was confirmed in proliferation experiments and functional assays. Specifically, amino acids and central carbon metabolites were attenuated while higher abundances were observed for fatty acids and most nucleic acid related metabolites. Interestingly, exposure to model xenoestrogens appeared to partially counteract these effects.

## Introduction

Breast cancer is the predominant type of cancer in women and accounts for approximately 25% of all cases in females. While the Surveillance, Epidemiology, and End Results (SEER) program of the US National Cancer Institute (NCI) recently reported a five-year relative survival rate of 88.8% for the years 2006-2012 (Jemal et al., 2017), the incidence of breast cancer continues to rise, with an overall increase from 105 to 130 per 100,000 population between 1975 and 2014 (NCI, 2017). Many risk factors for breast cancer are known to be hormone-mediated, including obesity, hormone replacement therapy during menopause, late pregnancies or nulliparity, and exposure to endocrine-disrupting xenoestrogens such as phyto-/mycoestrogens through diet or the environment (IARC, 2014; Yager and Davidson, 2006). More than 70% of breast cancer cases are classified as estrogen receptor (ER) positive (Howlader et al., 2014) and the manipulation of estrogen action has become a core aspect of breast cancer treatment (Musgrove & Sutherland, 2009).

To date, endocrine therapy is the most commonly administered first-line therapy for ER positive metastatic breast cancer (Nagaraj and Ma, 2015), because these cancer cells require estrogen supply for their growth (Yager and Davidson, 2006). The two receptor subtypes, ERα and ERβ, regulate gene expression through genomic and non-genomic signaling pathways in response to estrogen exposure affecting breast cancer cell growth and proliferation (Shanle and Xu, 2011). Selective estrogen receptor modulators (SERMs) may therefore be able to inhibit the estrogen receptor function (e.g. tamoxifen). Alternatively, aromatase (i.e. CYP19A1) inhibitors, such as anastrozole or letrozole (LET), block the initial production of estrogen. Another well-known general target in cancer therapy is cell cycle regulation and in recent years, three orally applied agents have been developed which selectively target the cyclin-dependent protein kinases (CDKs) 4 and 6, namely palbociclib (PAL), abemaciclib, and LEE011 (Mayer, 2015). PAL (Ibrance®, PD0332991) targets the adenosine triphosphate (ATP) binding site of CDK4-cyclin D and CDK6-cyclin D complexes and induces cell cycle arrest in the G(1) phase (Johnson et al., 2016b). It is applied orally in combination with either the aromatase inhibitor LET or the ER antagonist fulvestrant (FULV). This treatment in combination with LET is used as initial endocrine-based therapy in postmenopausal women with ER positive, human epidermal growth factor receptor 2 (HER2) negative metastatic breast cancer (Dhillon, 2015; Mechcatie, 2015).

An accelerated FDA approval for this combined therapy was granted in 2015. It was based on a randomized phase 2 study (PALOMA-1) of 165 postmenopausal women, resulting in a progression-free survival rate of about 20.2 months among the women treated with the combination of PAL and LET, compared with about 10.2 months among those treated with LET alone (Finn et al., 2015). The phase 3 study PALOMA-2 recently confirmed that the addition of PAL to the standard endocrine therapy (LET) significantly improved outcomes in the first-line treatment of breast cancer (Finn et al., 2016) making CDK4/6 inhibition in combination with antiestrogens the new standard for the treatment of advanced ER positive breast cancer (Wolff, 2016). Despite these groundbreaking clinical outcomes, the metabolic mechanisms underlying the drugs synergism at the metabolome level are yet unknown.

Considering the rise in breast cancer incidence, changes in diet have been considered as a potential cause. Phytoestrogens are plant-derived and food related compounds with known estrogenic activity. Epidemiological evidence has implied that diets rich in phytoestrogens reduce the incidence of breast cancer, hence they are applied as so-called ‘natural’ alternatives to hormone replacement therapy. However, their estrogenicity is also known to stimulate growth in experimental models of breast cancer (Rice and Whitehead, 2006). There are several classes of phytoestrogens with the most prominent being the isoflavones, such as genistein (GEN), and the lignans. Some fungi are also able to produce secondary metabolites with estrogenic activity; these are referred to as mycoestrogens. This includes the non-steroidal estrogenic mycotoxin zearalenone (ZEN) which is the most prominent representative of this class. A strong indication that ZEN and some of its metabolites play an important role in the promotion of hormone-dependent tumors has been previously reported, particularly for those arising from breast and endometrium (Dees et al., 1997; Pazaiti et al., 2012). ZEN and its reduced metabolites have been shown to interact with ERs and stimulate breast cancer cell growth in cell culture experiments (Khosrokhavar et al. 2009). Since LET’s mode of action is to block the production of endogenous estrogen and act as an aromatase inhibitor, it is relevant to evaluate if dietary xenoestrogens may interfere with LET and combined PAL+LET treatment.

Mass spectrometry-based metabolomic analysis allows for the comprehensive measurement of small molecules and their modulation upon perturbance in a biological system. Since metabolites are the end products of regulatory processes, they provide a functional readout of the cellular state and their levels can be regarded as the ultimate response of a system to genetic or environmental challenge (Patti et al., 2012b). Metabolomics can reveal biologically relevant alterations resulting from xenobiotic exposure or any other environmental challenge (Johnson et al., 2012). While this technology is routinely used for biomarker discovery, recent efforts are aiming towards deciphering mechanisms at the systems-level (Huan et al., 2017; Johnson et al., 2016a). Although it is the youngest of the major omics-disciplines, this field has made remarkable progress within the past decade mainly based on developments in analytical and bioinformatic technology (Johnson et al., 2015; Warth et al., 2017a).

The aim of this study was to elucidate the overall response of either individual PAL and LET dosing or a combination treatment on cell metabolism in ER positive breast cancer cells. We evaluated metabolomics data using XCMS Online and a systems biology/pathway prediction tool to provide measures for significantly altered metabolites and integrated metabolic pathway analysis. Moreover, we addressed the impact of dietary xenoestrogens on this combined treatment for the first time.

## Results and Discussion

### Metabolic effect of dosing individual agents

To gain a deeper understanding of the drugs effect on cellular metabolism, breast cancer cells were dosed with individual agents and a combination of PAL+LET (Figure 1). Meta-analyses were used for initial data evaluation to compare between the control cells, cells dosed with the single agents and cells receiving the combined dose (Patti et al., 2012a). As illustrated in Figure 2A, the individual drugs have exclusive effects on cellular metabolism with only nine statistically significant common metabolic features changed in both treatments. System-wide, exposure towards PAL at a concentration of 200 nM over a period of 48 h lead to a significant alteration of metabolic features (524) after filtering by p-value and fold change (see experimental section). Incubation with 10 nM LET resulted in a similar number of dysregulated features (459). These distinct effects of the agents is reflected by the pathway cloud plots shown in Figure S2, which report predicted modified metabolic pathways based on the recently implemented systems biology functionality within XCMS Online (Forsberg et al., 2017; Huan et al., 2017). This tool allows for the rapid prediction of dysregulated pathways without time consuming metabolite identification. While caution when interpreting these results is warranted, many of the key metabolites involved in the predicted pathways including several molecules associated with central carbon and nucleotide metabolism have been manually verified in a second step (Table 1). The drug concentrations were chosen to mimic plasma levels in patients undergoing therapy and based on previous reports (Finn et al. (2009), Johnson et al. in preparation).

**Fig. 1.**
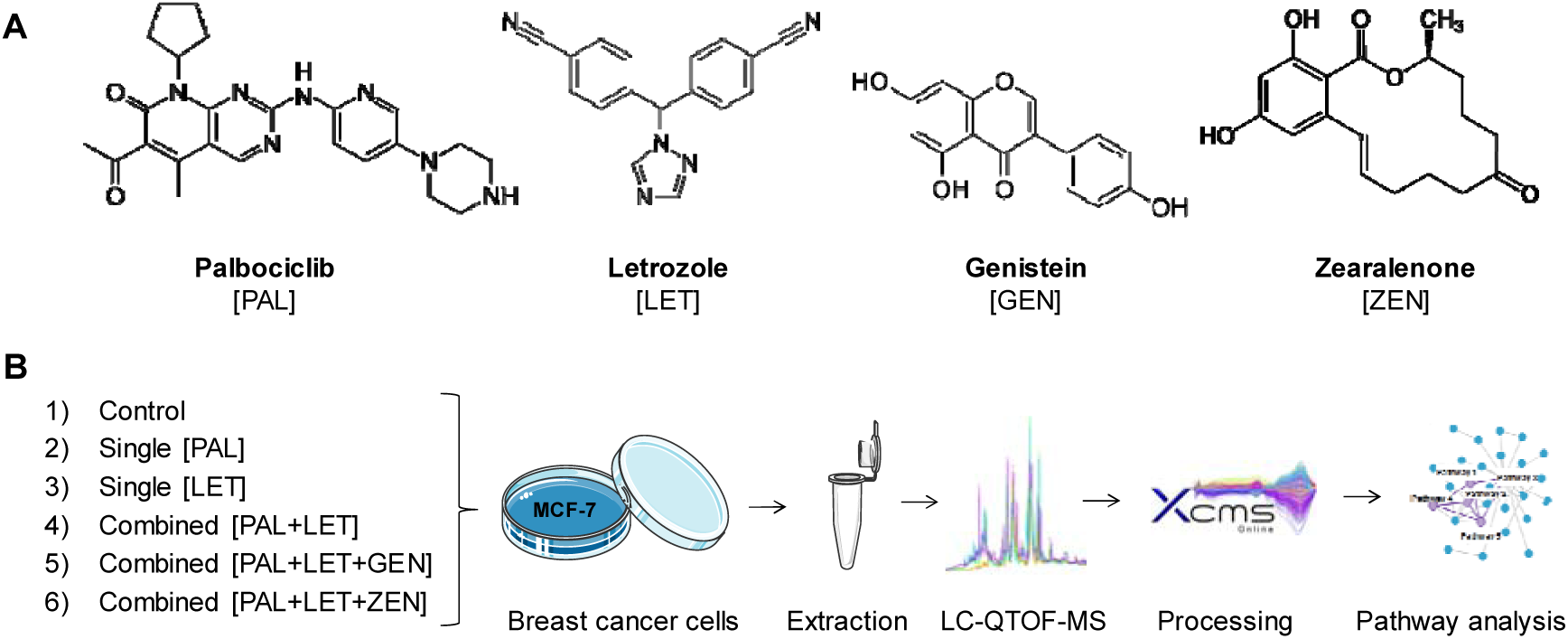
Overview of (A) the chemical structures of tested drugs and dietary xenoestrogens and (B) the experimental design of the applied global metabolomics approach. (A) The selective CDK 4/6 inhibitor palbociclib and the aromatase inhibitor letrozole are a first-line treatment for ER positive and HER negative advanced breast cancer. The isoflavone genistein and the mycotoxin zearalenone are dietary xenoestrogens exhibiting affinity towards estrogen receptors. (B) The untargeted metabolomics workflow was applied to the individual and the combined treatment of cancer drugs to study their synergism at the metabolomics level. In addition, the effect of the two xenoestrogens on the combined drug dosing was investigated.

**Fig. 2.**
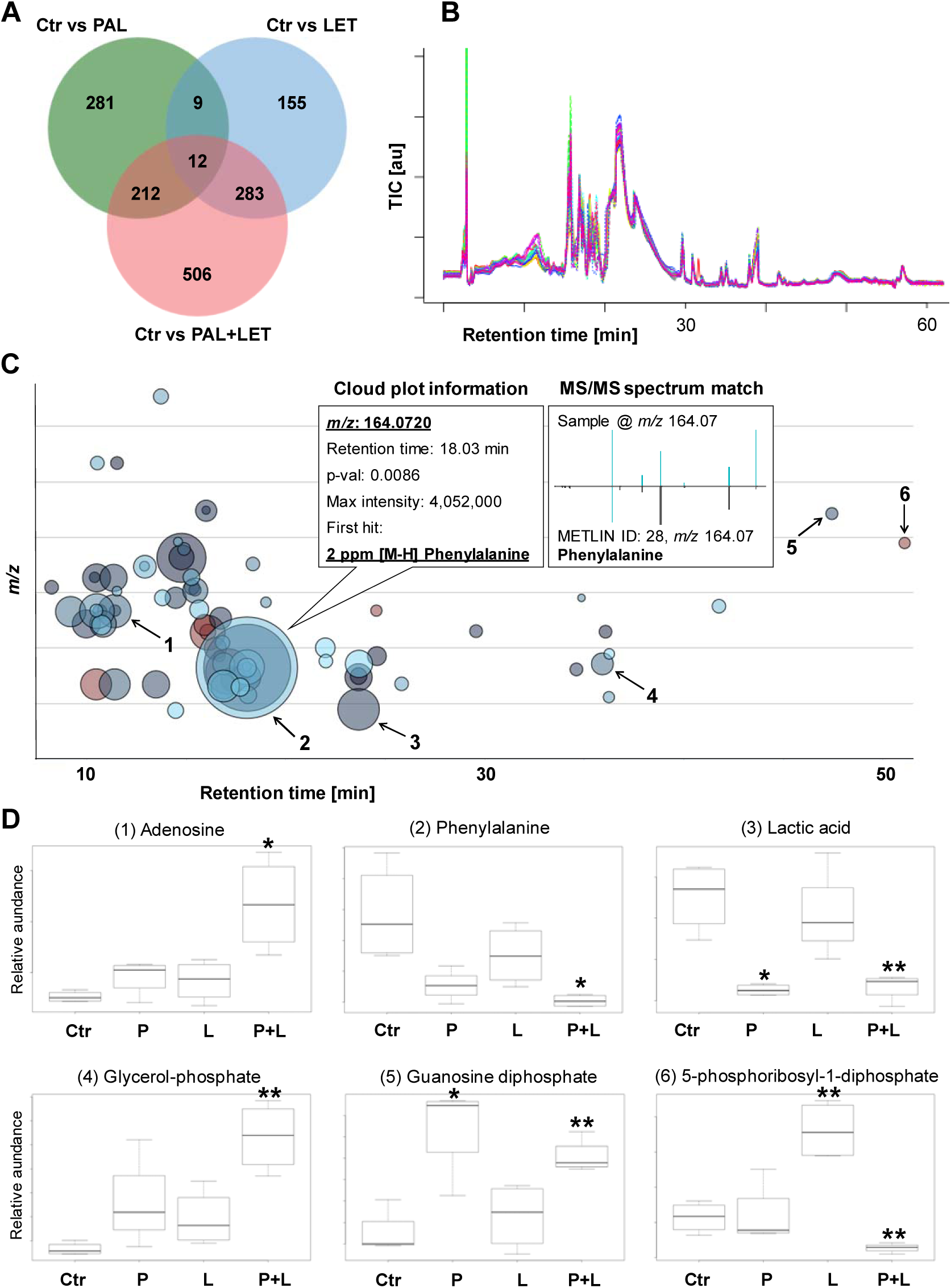
Comparison of single agent dosing versus the combined palbociclib + letrozole treatment in MCF-7 breast cancer cells. (A) Venn diagram of meta-analyses between the individual and the combined dosing. Combined exposure towards PAL and LET resulted in approximately twice as many altered metabolic features (1,013) than in the single treatments (PAL 524 and LET 459) after 48 h incubation (fold change >1.5, p-value <0.05). (B) Total ion chromatograms (TIC) of the HILIC-LC-MS measurements were automatically corrected for retention time shifts and improved the quality of the mass spectrometric data underlying (A), (B), and (C). The multi-group metabolomics cloud-plot shown in (C) and the boxplots (D) highlight specific changes of the cancer cells metabolome following drug dosing. Box plots (D) were extracted from (C) for six selected metabolites (1-6) whose abundances changed significantly in the combined dosing group. Palbociclib, letrozole, and the combination of both were compared to the solvent control (means ± SEM; n = 4; *p < 0.05, **p < 0.01, ***p < 0.001).

**Table 1.** Selected metabolites significantly dysregulated in MCF-7 cells after 48 h dosing with palbociclib (200 nM; n=4), letrozole (10 nM; n=4) or a combination of palbociclib and letrozole (200 nM and 10 nM, respectively; n=4). Values are fold changes when comparing dosed groups to control (dosed/control). Unpaired two-tailed t-test with 0.05 set as p-value for statistical significance. Trend is indicated for the combined treatment with + (1.0-1.5 fold), ++ (1.5-2.0 fold), +++ (> 2.0 fold), - (1.0- 0.75 fold), -- (0.75-0.5 fold), and --- (< 0.5 fold).

| Metabolite name | Palbociclib |  | Letrozole |  | Combination |  | Trend |
| --- | --- | --- | --- | --- | --- | --- | --- |
|  | Fold change | p-value | Fold change | p-value | Fold change | p-value |  |
| Central carbon metabolism |  |  |  |  |  |  |  |
| Glucose 6-phosphate | n.s. | n.s. | 1.5 | 0.0409 | 1.5 | 0.0086 |  |
| Glycerate 3-phosphate | 0.5 | 0.0114 | n.s. | n.s. | n.s. | n.s. |  |
| Phosphoenol pyruvate | n.s. | n.s. | 2.5 | 0.0094 | n.s. | n.s. |  |
| Ribose | 0.6 | 0.0389 | n.s. | n.s. | 0.3 | 0.0023 | --- |
| Ribose 5-phosphate | n.s. | n.s. | n.s. | n.s. | 8.6 | 0.0003 | +++ |
| Ribulose 1,5-diphosphate | n.s. | n.s. | 1.6 | 0.0018 | 0.5 | 0.0109 | --- |
| 5-phosphoribosyl-1-diphosphate | n.s. | n.s. | 2.2 | 0.0019 | 0.6 | 0.0088 | -- |
| 6-Phosphogluconic acid | 0.6 | 0.0178 | n.s. | n.s. | 0.6 | 0.0018 | -- |
| Lactic acid | 0.6 | 0.0133 | n.s. | n.s. | 0.6 | 0.0027 | -- |
| Malic acid | 0.8 | 0.0494 | n.s. | n.s. | 0.8 | 0.0103 | - |
| Nucleotide metabolism |  |  |  |  |  |  |  |
| Adenine | n.s. | n.s. | n.s. | n.s. | 6.4 | 0.0063 | +++ |
| Adenosine | n.s. | n.s. | n.s. | n.s. | 5.6 | 0.0131 | +++ |
| Guanosine | n.s. | n.s. | n.s. | n.s. | 2.2 | 0.0002 | +++ |
| Guanosine diphosphate | 1.7 | 0.0149 | n.s. | n.s. | 1.5 | 0.0015 | + |
| Guanosine triphosphate | 1.5 | 0.0352 | n.s. | n.s. | 1.5 | 0.0028 | + |
| Uridine | n.s. | n.s. | 2.4 | 0.0037 | 3.8 | 0.0011 | +++ |
| Deoxyuridine | 2.8 | 0.0478 | n.s. | n.s. | 6.0 | 0.0008 | +++ |
| Uridine monophosphate | 0.7 | 0.0309 | n.s. | n.s. | 0.4 | 0.0002 | --- |
| Amino acids |  |  |  |  |  |  |  |
| Phenylalanine | n.s. | n.s. | n.s. | n.s. | 0.5 | 0.0121 | -- |
| Leucine | n.s. | n.s. | n.s. | n.s. | 0.5 | 0.0295 | -- |
| Threonine | n.s. | n.s. | n.s. | n.s. | 0.7 | 0.0152 | -- |
| Lysine | n.s. | n.s. | n.s. | n.s. | 0.8 | 0.0103 | - |
| Aspartate | n.s. | n.s. | n.s. | n.s. | 0.7 | 0.0009 | -- |
| Glutamine | n.s. | n.s. | n.s. | n.s. | 0.7 | 0.0119 | -- |
| Glutamate | n.s. | n.s. | n.s. | n.s. | 0.8 | 0.0037 | - |
| Arginine | n.s. | n.s. | n.s. | n.s. | 0.7 | 0.0045 | -- |
| Fatty acids metabolism & biosynthesis |  |  |  |  |  |  |  |
| Caproic acid | 1.3 | 0.0061 | n.s. | n.s. | 2.5 | 0.0028 | +++ |
| Heptanoic acid | 1.2 | 0.0105 | n.s. | n.s. | 2.3 | 0.0010 | +++ |
| Caprylic acid | 1.2 | 0.0168 | 1.5 | 0.0001 | 2.1 | 0.0021 | +++ |
| Nonanoic acid | n.s. | n.s. | n.s. | n.s. | 1.7 | 0.0006 | ++ |
| Capric acid | n.s. | n.s. | n.s. | n.s. | 1.3 | 0.0042 | + |
| Undecanoic acid | 1.4 | 0.0038 | 1.2 | 0.0301 | 1.7 | 0.0005 | ++ |
| Lauric acid | 1.4 | 0.0198 | n.s. | n.s. | 1.6 | 0.0001 | ++ |
| Myristic acid | 1.6 | 0.0005 | n.s. | n.s. | 1.5 | 0.0018 | + |
| Pentadecylic acid | 1.6 | 0.0029 | 1.3 | 0.0283 | 1.8 | 0.0014 | ++ |
| Arachidonic Acid | 1.9 | 0.0215 | n.s. | n.s. | 1.8 | 0.0069 | ++ |
| Glycerol 3-phosphate | n.s. | n.s. | n.s. | n.s. | 2.8 | 0.0006 | +++ |
| 1-Palmitoyl Lysophosphatidic Acid | n.s. | n.s. | n.s. | n.s. | 3.4 | 0.0026 | +++ |
| 1-Oleoyl Lysophosphatidic Acid | n.s. | n.s. | n.s. | n.s. | 3.4 | 0.0058 | +++ |

n.s. not significant

While LET was shown to reduce proliferation of MCF-7 cells in the micromolar (Ismail et al., 2013; Rawat et al., 2013) and PAL even in the nanomolar range (Finn et al., 2009), our metabolomics results suggest no major effect on the proliferation of cancer cells. Typically, the stimulation of pathways related to glucose uptake, glycolysis, as well as lipid-, protein- and nucleotide synthesis is an indication of cellular proliferation (Agathocleous and Harris, 2013). However, given the very low dose of LET (10 nM) and the short incubation time of 48 h compared to other studies (Finn et al. (2009), Johnson et al. in preparation), this was not unexpected. In the single treatments, no effect on amino acid metabolism and only a minor response on nucleotide metabolism was observed for both drugs. However, PAL alone reduced several intermediates of the central carbon metabolism (glycerate-3-P, ribose, 6-phosphogluconic acid, lactic acid, malic acid) while for others (glucose-6-phosphate, ribulose-1,5-diphosphate, 5-phosphoribosyl-diphosphate) a slight increase was seen in the LET group. Only a few fatty acids were upregulated by LET with a generally more potent effect of PAL (Table 1).

### Combined agents potentiate the effects on cellular metabolism

The meta-analyses performed on XCMS Online illustrated the overall highly pronounced impact of the combination treatment compared to individual exposures by way of a Venn diagram, together with a metabolomics cloud plot and extracted box- and whisker plots (Figure 2A). The combination of PAL and LET resulted in 1,013 altered metabolic features in the breast cancer cells. Many of these metabolic features, however, may result from the same molecule due to the formation of adducts or in-source fragment ions (Zamboni et al., 2015). From this data set more than 100 significantly dysregulated metabolites (fold change >1.5, p-value <0.05) were derived and 58 metabolites were putatively identified. Many of those were matched to key molecular building blocks with a prominent role in cellular metabolism. Table 1 reports on the relative changes upon single and combined treatments for major metabolites associated with central carbon, nucleotide, amino acid, and fatty acid metabolism. In line with the meta-analysis (Figure 2), the effects on cellular metabolism following combined treatment is enhanced and draws a distinct picture: On the one hand, all significantly altered amino acids and central carbon metabolites (except ribose-5-phosphate) were clearly less abundant after PAL+LET dosing. On the other hand, fatty acids as well as nucleobases, nucleosides and nucleotides (except UMP) showed the opposite behavior with generally higher abundances (Table 1 and Figure 2). This behavior is corroborated by the results of the metabolic pathway prediction (Figure S2, Huan et al. (2017)). We speculate that the increased levels of nucleic acid precursors might be caused by the PAL mediated inhibition of CDK 4/6 and the resulting cell cycle arrest. This mode of action would prevent the utilization of precursors in the S phase where high concentrations are required for DNA replication (Lane and Fan, 2015).

The lower abundance of amino acids suggests that the mammalian target of rapamycin (mTOR) pathway, which is a negative target of AMP activated protein kinase (AMPK), is attenuated (Agathocleous and Harris, 2013). When activated by high amino acid levels, this pathway typically promotes anabolic reactions with multiple roles in cell differentiation and a stimulation of glycolysis which was not apparent in our data. Since basically all amino acids were less abundant in the combined dosing group it appears reasonable to speculate that a master regulator, such as mTOR, is involved in the observed molecular events. mTOR acts as a downstream effector for many oncogenic pathways and deregulation of mTOR-signaling is a hallmark of many human cancers (Saxton and Sabatini, 2017).

To investigate if the combined PAL+LET treatment affects the mTOR pathway, we set out to determine the phosphorylation state of the mTOR downstream targets p70-S6-kinase and ribosomal protein S6. PAL+LET reduced phosphorylation of the proteins in the two tested ER positive cell lines MCF-7 and T47D, indicating an inhibitory effect of the combined treatment on activation of S6 kinase by mTOR (Figure 3).

**Fig. 3.**
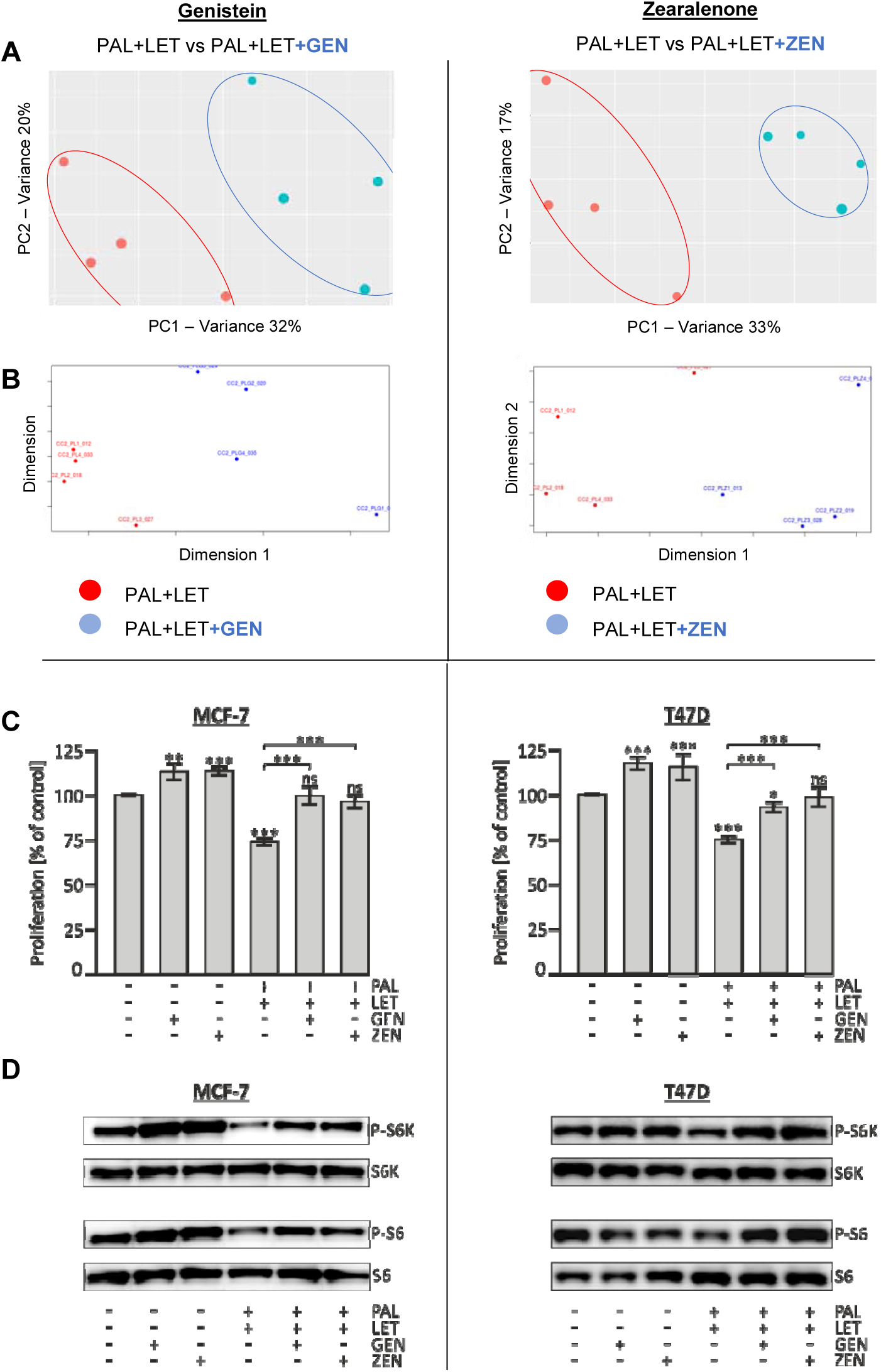
Dietary xenoestrogens counteract the metabolic and anti-oncogenic effects of PAL+LET in human breast cancer cells. The combined treatment (PAL+LET) was compared to the combined treatment plus either 1 µM genistein (GEN) or 100 nM zeralenone (ZEN). (A) PCA score plots: The first (PC1) and second (PC2) principal component explain 32% and 20% of the variance respectively for the experiment containing GEN, with very similar variances for ZEN (33% and 17%). (B) Non-metric multidimensional scaling visualizes the similarity level of individual samples in a dataset and additionally suggests the distinct effect of the xenoestrogens on the combined PAL+LET dosing samples. (C) The effect of GEN and ZEN on the PAL+LET treatment was analyzed in cell proliferation assays. The human breast cancer cell lines MCF-7 and T47D were exposed to indicated drugs and/or xenoestrogens for 72 h and viable cells were quantified using the resazurin-assay. Shown are the percentages of individual samples relative to the control sample (DMSO only) which was set to 100% (means ± SD; n = 4; *p < 0.05, **p < 0.01, ***p < 0.001). (D) Impact of xenoestrogens and/or combined PAL+LET treatment on phosphorylation state of mTOR downstream targets. Cells were exposed to the indicated compounds for 48 h. Western blot analysis of phospho-p70-S6-kinase (P-S6K), total S6K, phospho-S6 (P-S6), and total ribosomal protein S6. The images are representative of three independent experiments.

Interestingly and in contrast to a recent study on the combined effects of PAL and the ER antagonist FULV (Johnson et al., in preparation), no metabolic signatures of cell death were observed in the study at hand. Most notable, nucleotide metabolism was overall upregulated after two days of PAL+LET exposure while PAL+FULV resulted in some metabolites with dramatically decreased abundances (e.g. inosine monophosphate was downregulated 16-fold). Ribose-5-phosphate (R5P) was the only metabolite derived from the central energy metabolism not attenuated but instead significantly upregulated (8-fold, p-value 0.0003). R5P is a product but also an intermediate of the pentose phosphate pathway and behaved again differentially than in the cited PAL+FULV paper where decreased concentrations were reported (Johnson et al., in preparation). Thus, FULV and LET appear to have different effects on cellular metabolism when combined with PAL, and it remains to be determined whether the metabolic consequences of these drug treatments are responsible for the improved clinical outcome seen in patients on combination therapies (Cristofanilli et al., 2016; Finn et al., 2016).

### Role of xenoestrogens during combined agent dosing

Pair-wise and multigroup analysis within XCMS Online were utilized to evaluate the effect of the phytoestrogen GEN and the mycoestrogen ZEN on the combined PAL+LET treatment. Principal component analysis (PCA) and non-metric multidimensional scaling (MDS) of the obtained data allowed for an unbiased assessment and visualization of their impact on a global scale (Figure 3 A-B). Using these non-supervised, multivariate analyses, it became apparent that the investigated xenoestrogens counteracted the metabolic effects of the combined PAL+LET treatment group. This behavior was confirmed for specific metabolites which seemed to be particularly interesting players in the combined drug action (Table 1). Most of the highly dysregulated features observed in the univariate data were also found in the PCA model. The antagonistic effect of xenoestrogens on the combined drug treatment was also confirmed in cell proliferation assays using the ER positive cell lines MCF-7 and T47D. Proliferation of both cell lines was markedly reduced after exposure to PAL+LET, when compared to untreated cells. Strikingly, co-administration of PAL+LET with either of the two xenoestrogens GEN or ZEN restored cell proliferation to levels that were comparable to non-treated cells (Figure 3C). This functional assay together with our metabolomics data provides evidence that dietary xenoestrogens have the potential to circumvent the anti-oncogenic effects of the PAL+LET combination therapy.

It has been reported in breast epithelial cells that the xenoestrogen bisphenol A induces activation of the mTOR pathway and leads to marked resistance to the mTOR-inhibitor rapamycin (Goodson et al., 2011). To investigate if the observed effects of GEN and ZEN on the combined PAL+LET treatment also affect the mTOR pathway, the phosphorylation state of the mTOR downstream targets p70-S6-kinase and ribosomal protein S6 were determined after xenostrogen co-exposure as well. Our results show that the inhibitory effect of the combined drug treatment is antagonized by the tested xenoestrogens GEN and ZEN (Figure 3 D) which is in line with the obtained metabolite data. mTOR activity is negatively regulated by low intracellular amino acid levels (Agathocleous and Harris, 2013). In MCF-7 cells PAL+LET significantly reduced the abundancy of numerous metabolites, notably many amino acids. However, addition of GEN or ZEN to the drug combination counteracted this inhibitory effect of PAL+LET and led to cellular amino acid levels that were comparable to untreated cells in many cases (Figure 4).

**Fig. 4.**
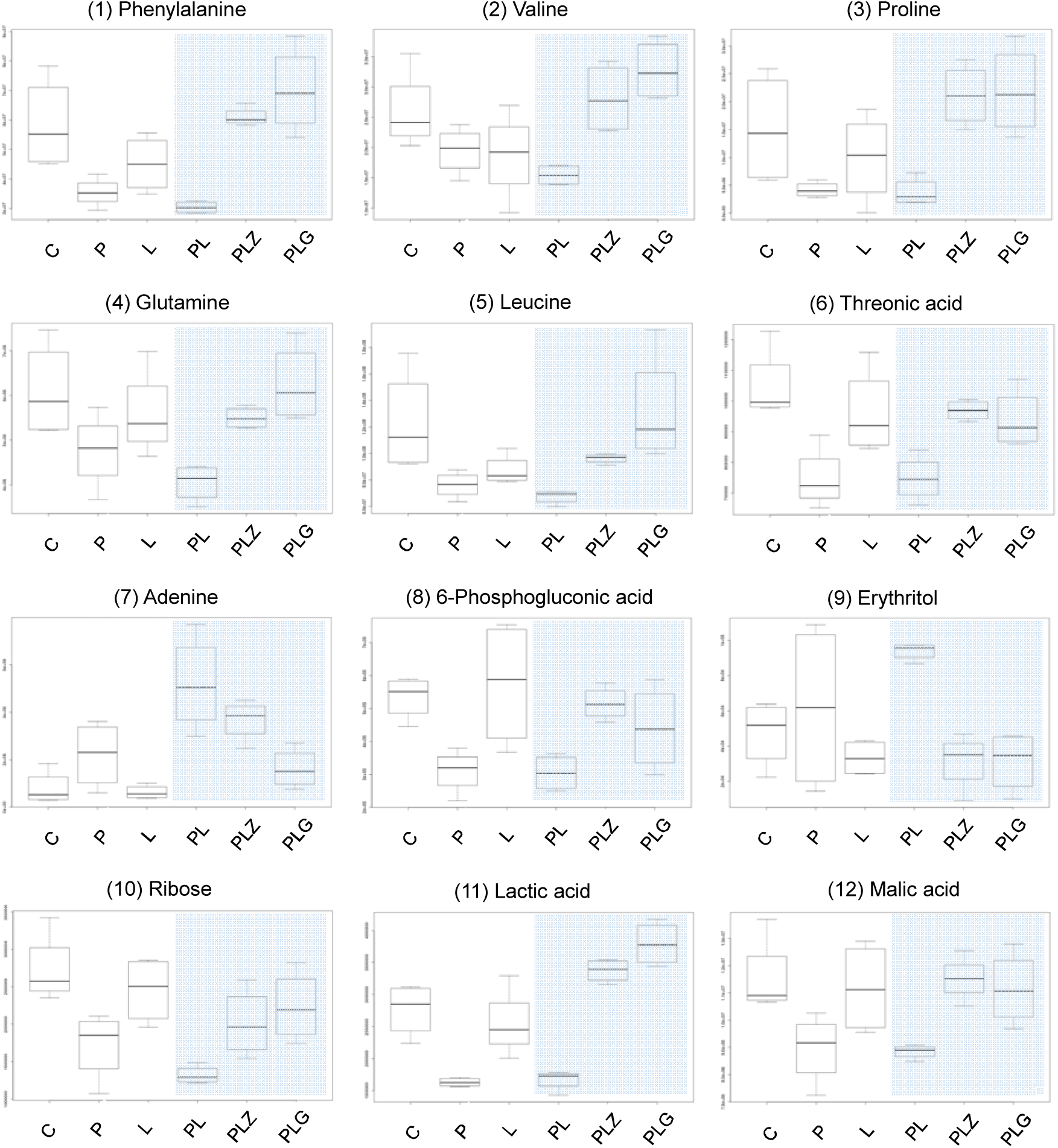
Relative abundances of selected key metabolites identified during combined PAL+LET (PL) dosing were significantly altered by the dietary xenoestrogens zearalenone (PLZ) and genistein (PLG). The effect of the two food estrogens (100 nM ZEN or 1 µM GEN) added to the combined treatment was compared to the control, the single agents (PAL, P; LET, L), and the combined treatment (n=4). The impact of the estrogens on the combination is highlighted by the blue shade. The y-axis represents relative metabolite abundances [a.u.].

Breast cancer is a multi-faceted disease with dietary factors and hormones being key elements in its development. Diet has long been acknowledged for its role to contribute to cancer risk, however, the molecular foundations of this phenomenon remain poorly understood (Sullivan and Vander Heiden, 2017). Bioactive food constituents and contaminants have been associated with both beneficial and potentially adverse effects on cancer susceptibility, progression, and outcome in general. In particular phytochemicals such as polyphenols have been used for the treatment of cancer, however, despite recent advances empowered by new metabolomic technology, their effect on cancer metabolism has not yet been fully clarified (Brasili and Filho, 2017). This is especially true for the impact of specific compounds during chemotherapy. There is considerable epidemiological evidence from Asian countries that high soy diets (i.e. rich in isoflavones such as GEN) lower the incidence of breast cancer (Hilakivi-Clarke et al., 1999). Typically, Asian women consume substantially more soy food than women in Western-style populations resulting in high phytoestrogen early-life exposures. Thus, it was suggested that the time point in life when exposure begins seems to be a critical parameter for a potential protective effect (Adlercreutz, 2002; Blei et al., 2015). A recent multiethnic US study associated higher dietary isoflavone intake with lower mortality in women with HR negative tumors and those not treated with hormone therapy while no negative impact on all-cause mortality was found for patients receiving (self-reported) hormone therapy (Zhang et al., 2017). Nechuta et al. (2012) described an association between higher consumption of soy-based foodstuff following a breast cancer diagnosis with improved treatment outcomes and reduced recurrence rates in more than 9,500 survivors from China and the US. Interestingly, isoflavone intake was inversely associated with recurrence among both groups despite the large differences in dietary soy consumption by country.

Contradictory to these epidemiological surveys, GEN as well as other phytoestrogens stimulated the growth of ER+ cell lines in vitro (Rice and Whitehead, 2006). This paradox might be caused by the stronger affinity of phytoestrogens towards the ERβ than the ERα form of the estrogen receptor (Rice and Whitehead, 2006). The xenoestrogens GEN and ZEN trigger a similar response on a significant share of metabolites in the treated cells in this study (Figure S3) even though GEN has a higher affinity towards the ERβ than the ERα form of the receptor (Rice & Whitehead, 2006) and ZEN interacts with both in a comparable manner (Takemura et al., 2007). Activation of ERα is associated with growth promotion in breast tumors, while the role of ERβ is less well understood. In general, affinity of phytoestrogens to the ERs is much lower than that of the principal endogenous estrogen 17β-estradiol.

Other groups have previously described changes in metabolite concentrations in breast cancer cells or xenograft models upon incubation with phytohormones. For example Jäger et al. (2011) found increased levels of amino acids and arachidonic acid in two breast cancer cell lines upon high resveratrol doses. Ju et al. (2008) showed that dietary GEN can reverse the inhibitory effect of LET on tumor growth and adversely impact breast cancer therapy. The authors even concluded that caution is warranted for consumption of dietary GEN by postmenopausal women with estrogen-dependent breast cancer taking LET treatment (Ju et al., 2008). Given the results obtained in this in vitro study it seems not unlikely that the phytoestrogen GEN and the mycoestrogen ZEN may also influence the described PAL+LET combination therapy through their bioactivity in an in vivo scenario. As we aimed to unravel the effects on metabolism on baseline xenoestrogen concentrations realistic in a Western style society (Soukup et al., 2016; Verkasalo et al., 2007; Warth et al., 2013), it is important to note that the described alterations of metabolism, cellular proliferation, and mTOR signaling were caused by rather low concentrations. It is possible that individuals consuming high soy (GEN) or high cereal (ZEN) diets may exceed these concentrations in their plasma.

The specific reverse action of xenoestrogens on proliferation, oncogenic signaling (Figure 3), and multiple key metabolites of the combined PAL+LET dosing (Figure 4) raises the question whether these dietary estrogens might have the potential to antagonize the synergistic effect of the combined treatment in breast cancer patients which led to its accelerated FDA approval (Finn et al., 2015). To ultimately determine if ZEN and/or GEN constitute a potential dietary risk factors for the progression of hormone dependent cancers or their treatment, further in vivo evidence is essential. To accomplish this task, it will be crucial to link sensitive exposure assessment of dietary estrogens to patient metabolism and therapy outcomes. These outstanding challenges are well-aligned with the increasing interest in assessing complex drug-exposome interactions utilizing global multi-omics and systems toxicology approaches. Innovative metabolomics technology was proposed to become a key driver in this new research arena at the edge of disease progression/treatment, environmental exposures and nutrition/food research (Beger and Flynn, 2016; Jones, 2016), a perspective supported by the data presented in this work.

## Conclusions & Outlook

We describe the metabolic response of breast cancer cells on combined PAL and LET treatment and demonstrate their powerful impact on cellular metabolism, while the drugs dosed individually caused only minor effects. These results are in line with a phase 2 clinical trial of this combination treatment, the outcome of which led to its accelerated FDA approval (Finn et al., 2015). Untargeted metabolomics helped to describe the overall cellular response, and it demonstrated that the pathway prediction tool expedited and refined data processing and evaluation. The dietary xenoestrogens GEN and ZEN showed a distinct effect on cellular metabolism, oncogenic signaling, and cell proliferation during the combined treatment. Our results shed light on the vast impact bioactive food-related molecules may pose on cancer metabolism and treatment. It will be important to translate the results obtained in these experiments to animal models and ultimately to patients to enable targeted nutritional recommendations for breast cancer patients while undergoing treatment.

## Significance

Breast cancer is the leading type of cancer in women and more than 70% are classified as the subtype hormone receptor (estrogen and progesterone) positive and human epidermal receptor (HER2) negative. Exposure to xenoestrogens has been suggested as a risk factor in these cancers. They might act as endocrine disruptors and affect hormone signaling leading to cancer and may even impact therapeutic response, especially regarding drugs interacting with the ER. For patients with ER positive, HER2 negative tumors, recent clinical evidence proved superior outcomes in those receiving a combination therapy of a cyclin dependent kinase 4/6 (CDK4/6) inhibitor (palbociclib, Ibrance®) in addition to the current standard endocrine therapy (letrozole). However, neither the underlying metabolic effects causing this synergistic combinatory effect, nor those on the individual agents have been investigated in an untargeted manner in cell models to date. Our global metabolomics data draws a distinct picture and describes effects of this combinatory treatment on the MCF-7 metabolome for the first time. Furthermore, we highlight the effects bioactive food chemicals may inflict on this novel therapy. Interestingly and potentially of high nutritional and clinical relevance, exposure to the tested model xenoestrogens appeared to counteract the metabolic and anti-proliferative effects of the combined therapy in vitro. Our study provides evidence that these food constituents/contaminants may attenuate the synergistic effect of the combined treatment in breast cancer patients.

## Footnotes

**Author Contributions** B.W. designed the research idea and experiments. B.W., P.R., and A.G. performed cell culture and mass spectrometric experiments. B.W., P.R., T.H., M.F., E.F., P.B., L.G., C.H.J., and G.S. analyzed and evaluated the data. All authors contributed to manuscript writing.

## Acknowledgements

The authors would like to express their sincere gratitude for critical and fruitful discussions towards J. Rafael Montenegro-Burke, Bill Webb, Catherine Bell(TSRI), Steven Dann (Pfizer Inc., La Jolla), Giorgia Del Favero and Doris Marko (University of Vienna). Minerva Tran and Linh Hoang (TSRI) are greatly acknowledged for skillful lab support. The authors further thank Steven Dann for providing PAL stock solutions and the Austrian Science Fund (FWF; Schrödinger fellowship J-3808 awarded to B.W.), the George E. Hewitt Foundation for Medical Research (P.R.), and the National Institutes of Health grants R01 GMH4368 and PO1 A1043376-02S1 for financial support.

## Data and Software availability

The datasets described in this work will be made available via the ‘XCMS Public’ data repository (<u>https://xcmsonline.scripps.edu/landing_page.php?pgcontent=listPublicShares</u>) once acceptance for publication.

## STAR METHODS

### CONTACT FOR REAGENT AND RESOURCE SHARING

Further information and requests for reagents may be directed to, and will be fulfilled by, the corresponding author Benedikt Warth

### EXPERIMENTAL MODEL DETAILS

#### Cell Lines

MCF-7 cells were obtained from ATCC (Manassas, VA) and cultured in Dulbecco’s Modified Eagle Medium F-12 Nutrient Mixture (Gibco-Life Tech, Grand Island, NY) without phenol red and supplemented with 10% fetal bovine serum (Sigma, St. Louis, MO) and penicillin-streptomycin (50 U/mL, Gibco-Life Tech, Grand Island, NY) at 37 C and 5% CO_2_. T47D cells were obtained from ATCC (Manassas, VA) and cultured in RPMI 1640 (Gibco-Life Tech, Grand Island, NY) supplemented with 10% FBS, 50 U/ml penicillin-streptomycin, and 0.2 U/ml bovine insulin.

### METHOD DETAILS

#### Cell culture

MCF-7 breast cancer cells, the most popular cell model to study ER positive breast cancers due to its exquisite hormone sensitivity (Holliday and Speirs, 2011), were used in all experiments whereas T47D cells were only examined in the proliferation and functional assays for confirmatory purposeFor metabolomics experiments, FBS was exchanged by charcoal-dextran stripped FBS, which provides low levels of hormones. Cells were routinely split and maintained in T-75 flasks at a confluence between 75-85%. For sub-culturing the cells were washed with PBS, detached with trypsin/EDTA and re-suspended in fresh culture medium. For the experiments cells were seeded in 6-well plates containing 3 ml cell culture medium. After 48 h cells were either incubated with common medium as control, PAL (200 nM), LET (10 nM), a PAL+LET combination (200 nM, 10 nM), or the combined treatment plus the phytoestrogen GEN (1 µM) or the mycoestrogen ZEN (100 nM) for 48 h. The drug concentrations were based on a realistic ratio and previous reports (Finn et al. (2009), Johnson et al. in preparation). For xenoestrogens, realistic concentrations were also in line with published data on food consumption/contamination and background concentrations in bio-fluids (Vejdovszky et al., 2017; Warth et al., 2013) were chosen to mimic a scenario as realistic as possible. For the duration of T47D experiments, phenol red-free RPMI 1640 and charcoal-dextran stripped FBS were used as well. Cell proliferation experiments were carried out in 96-well plates. 24 h after seeding (3,000 cells per well, 100 µl culture medium) drugs and/or xenoestrogens (200 nM PAL, 20 nM LET, 1 µM GEN, 100nM ZEN) were added and cells were incubated for 72 h. A 100x stock solution (1 mg/ml) of the redox dye resazurin was added to the cell culture medium to a final concentration of 10 µg/ml and after incubation at 37 C for 90 min, fluorescence intensities were quantified on a SYNERGY 4 instrument from BioTek (Winooski, VT). For phosphorylation assays cells were grown in 6-well plates and 48 h after drug and/or xenoestrogen (200 nM PAL, 20 nM LET, 1 µM GEN, 100 nM ZEN) exposure cell extracts were prepared (lysis buffer: PBS containing 0.5% Triton-X100, 1x protease inhibitor cocktail, 1 mM PMSF, 1 mM orthovanadate, and 50 mM sodium fluride). Cell lysates ware centrifuged at 15,000xg for 30 min at 4 C and protein concentration of the soluble fraction was determined by BCA assay. Lysates (15 µg total protein per sample) were subjected to SDS-PAGE and electroblotting onto PVDF membranes. Immunoblotting experiments with indicated antibodies were carried out as recommended by the supplier (Cell Signaling Technology).

Cell culture media and supplements were purchased from ATCC (Manassas, VA) and Fisher Scientific (Pittsburgh, PA). Pierce BCA protein assay kit was purchased from ThermoFisher Scientific (Waltham, MA). Antibodies (phospho-p70-S6-kinase, Thr389 #9205; total p70-S6-kinase #9202; phospho-S6 ribosomal protein, Ser235/236 #2211; total S6 ribosomal protein #2217) were purchased from Cell Signaling Technology (Danvers, MA).

### Metabolomics Protocol

#### Sample preparation

To extract the cells for global metabolomcis experiments, the 6-well plates were placed on ice and the medium was removed using a vacuum pump. Cells were washed twice with 1 mL of ice cold PBS. Then, 1 mL ice cold quenching solution (MeOH:ACN:H2O (2:2:1, v/v) was added and the cells were detached using a cell scraper and the cell suspension was transferred to 1.5 mL microcentrifuge tubes (Eppendorf, Hauppauge, NY). Samples were vortexed followed by three cycles of shock-freezing in liquid nitrogen and subsequent thawing at room temperature and sonication 4°C for 10 min. To precipitate proteins samples were incubated for 1 h at −20°C and centrifuged 15 min at 13,000 rpm and 4°C. The supernatant was evaporated to dryness in a vacuum concentrator (Labconco, Kansas City, MO) and the dried extracts reconstituted in ACN:H2O (1:1, v/v) according to their protein content as determined by the BCA assay. Following sonication for 10 min, and centrifugation (15 min, 13,000 rpm, 4°C) the supernatants were transferred to LC vials and stored at - 80°C until analysis. All experiments were performed in at least four biological repetitions.

#### LC-MS/MS instrumentation

Analyses were performed using a high performance liquid chromatography (HPLC) system (1200 series, Agilent Technologies) coupled to a Bruker Impact II quadrupole time-of-flight (Q-TOF) high resolution mass spectrometer (HR-MS; Bruker Daltonics, Billerica, MA). Cell extracts (4 µL) were injected onto a Luna aminopropyl, 3 µm, 150 mm × 1.0 mm I.D. column (Phenomenex, Torrance, CA) for hydrophilic interaction liquid chromatography (HILIC) analysis in ESI negative mode. HILIC was chosen to analyze predominantly polar metabolites which typically retain better than on reversed phase columns. The mobile phase was A 20 mM ammonium acetate and 40 mM ammonium hydroxide in 95% water and 5% acetonitrile and B 95% acetonitrile, 5% water. A linear gradient from 100% B (0-5 min) to 100% A (50-60 min) was applied at a flow rate of 50 µL/min. Acetonitrile (ACN; Fisher Scientific, Pittsburgh, PA), methanol (MeOH; Honeywell, Morris Plains, NJ) and water (J.T. Baker, Center Valley, PA) were all LC-MS grade. ESI source conditions were set as follows: negative polarity, gas temperature 220°C, drying gas (nitrogen) 6 L/min, nebulizer 1.6 bar, capillary voltage 4500V. The instrument was set to acquire over a m/z range from 50-1000 with the MS acquisition rate of 2 Hz. To ensure column re-equilibration and maintain reproducibility, a 12 min post-run was applied. For the acquisition of MS/MS spectra of selected precursors the default isolation width was set to 2 Da with MS and MS/MS acquisition rates of 4 Hz. For the generation of MS/MS spectra, 8 µL of the cell extracts were injected. The collision energy was set to 20-50 eV over the scan range.

### QUANTIFICATION AND STATISTICAL ANALYSIS

#### Processing and Statistical Analysis of Metabolomics Data

Data was processed using XCMS Online (Tautenhahn et al., 2012) with an α=0.05. Pairwise, multigroup and meta XCMS jobs were run to evaluate the acquired HR-MS files in the most comprehensive way. Data was displayed as a feature table and plotted as a cloud plot for both, the metabolomic and pathway cloud plots (Huan et al., 2017). These contained m/z and retention time information (where applicable), integrated intensities, observed fold changes across the sample groups, and statistical significance for each sample. Metabolites were relatively quantified based on their abundances. Bruker Compass Data Analysis 4.3 software was used for additional manual data evaluation and verification. Metabolite identification was based on matching MS/MS spectra, mainly through the METLIN database (Smith et al., 2005), which housed nearly one million unique metabolites at that time (Warth et al., 2017b). In many cases the metabolite identities were further validated by comparison with authentic reference standards.

#### Processing and Statistical Analysis of proliferation assays

Data obtained from four independent cell proliferation assays (four replicates each) were combined for statistical analysis and displayed in bar graphs. Statistical significance of individual samples compared to control were determined using unpaired two-tailed t-test with 0.05 set as p-value for significance.

### DATA AND SOFTWARE AVAILABILITY

The raw metabolomics data will be provide via XCMS Public. Metabolomics data was processed via XCMS Online which is a free cloud-based processing platform.

### KEY RESOURCES TABLE

| REAGENT or RESOURCE | SOURCE | IDENTIFIER |
| --- | --- | --- |
| **Antibodies:** |  |  |
| Phospho-p70 S6 Kinase (Thr389) Antibody | Cell Signaling Technology | Cat#9205S |
| Phospho-S6 Ribosomal Protein (Ser235/236) Antibody | Cell Signaling Technology | Cat#2211S |
| p70 S6 Kinase Antibody | Cell Signaling Technology | Cat#9202S |
| S6 Ribosomal Protein (5G10) Rabbit mAb | Cell Signaling Technology | Cat#2217S |
| **Chemicals, Peptides, and Recombinant Proteins** |  |  |
| Genistein | Sigma-Aldrich | G6649, CAS Number 446-72-0 |
| Zearalenone | Sigma-Aldrich | Z2125, CAS Number 17924-92-4 |
| Palbociclib (metabolomics experiments) | Pfizer Inc. | - |
| Palbociclib (proliferation and mTOR assays) | Sigma-Aldrich | Cat#PZ0199 |
| Sodium orthovanadate | Sigma-Aldrich | Cat#S6508 |
| Sodium fluoride | Sigma-Aldrich | Cat#S7920 |
| EDTA-free protease inhibitor cocktail | Sigma-Aldrich | Cat#11873580001 |
| Phenylmethanesulfonyl fluoride (PMSF) | Sigma-Aldrich | Cat#P7626 |
| Resazurin sodium salt | Sigma-Aldrich | Cat#R7017 |
| **Critical Commercial Assays** |  |  |
| BCA assay |  |  |
| Pierce BCA Protein Assay Reagent A | Thermo Scientific | Cat#23223 |
| Pierce BCA Protein Assay Reagent B | Thermo Scientific | Cat#23224 |
| **Deposited Data** |  |  |
| Metabolomics pairwise job_ C vs L | XCMS Online | #1133412 |
| Metabolomics pairwise job_ C vs P | XCMS Online | #1133413 |
| Metabolomics pairwise_ C vs PL | XCMS Online | #1133414 |
| Metabolomics multi-group job_ C vs P vs L vs PL | XCMS Online | #1123819 |
| Metabolomics pairwise job_PL vs PLZ | XCMS Online | #1123822 |
| Metabolomics pairwise job_PL vs PLG | XCMS Online | #1123823 |
| Metabolomics multi-group job_ C vs P vs L vs PL vs PLZ vs PLG | XCMS Online | #1123826 |
| **Experimental Models: Cell Lines** |  |  |
| MCF-7 | ATCC | ATCC® HTB-22™ |
| T47D | ATCC | ATCC® HTB-133™ |
| **Software and Algorithms** |  |  |
| XCMS Online | (Tautenhahn et al., 2012) | http://xcmsonline.scripps.edu |

## Supplemental Information

**Fig. S1.**
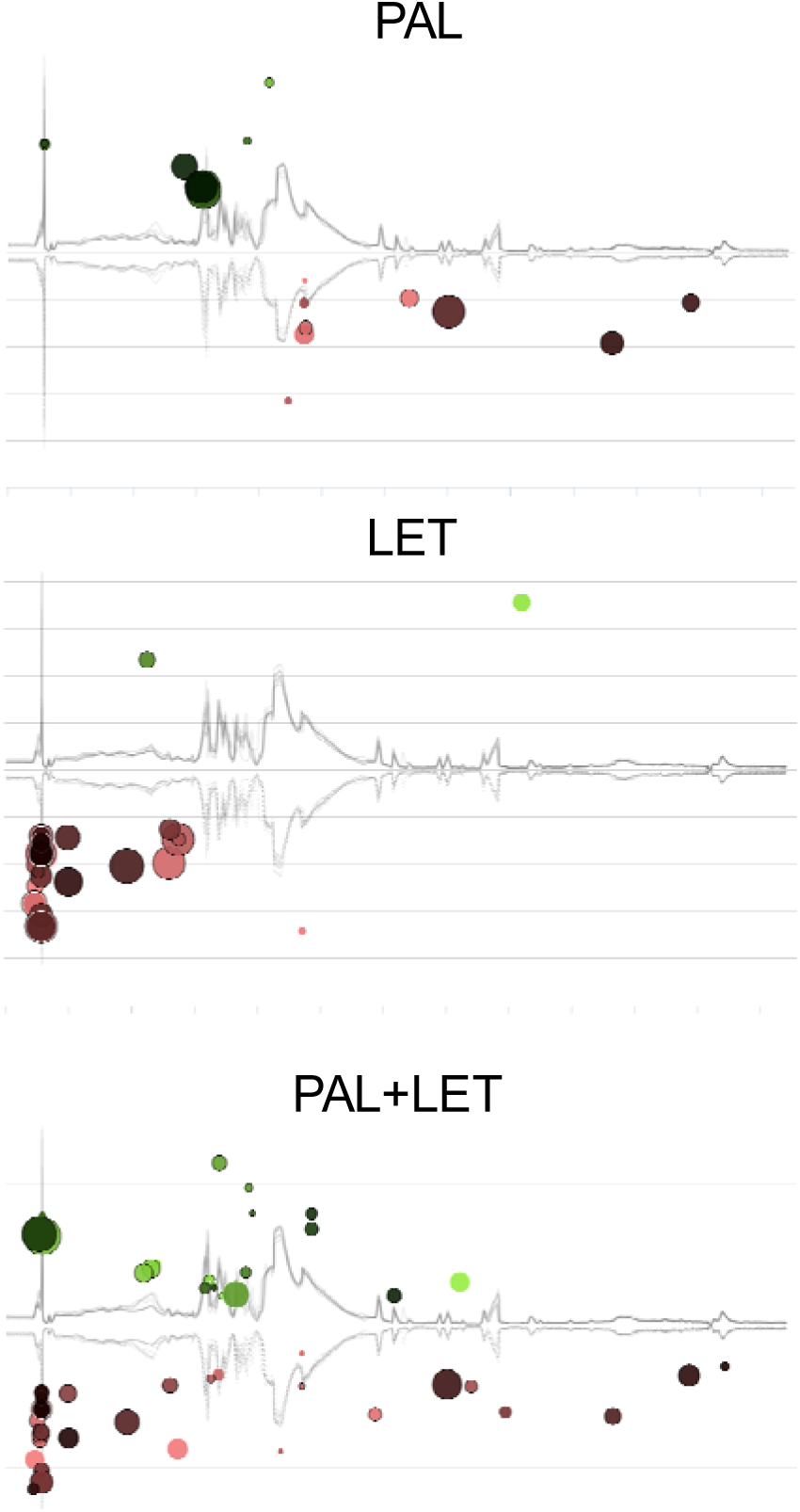
Pair-wise metabolomics cloud-plot analysis between the control and either PAL, LET, or PAL+LET as obtained for HILIC-LC-ESI-neg-MS measurements in MCF-7 treated cells. Palbociclib (200 nM), letrozole (10 nM), and the combination of both were compared to the vehicle (control) after 48 h of drug exposure to show the overall response of the metabolome following the individual or combined treatments. Following the combined treatment, there are clearly more metabolic features altered than following the individual treatments. Cloud plots show all highly significantly dysregulated metabolites (fold change > 1.5, p-value < 0.001). Metabolic features whose intensity were increased after treatment are shown on the upper part of the plot as green circles and those with decreases intensities are shown on the bottom as red circles. Bigger and brighter circles (features) correspond to larger fold changes and lower p-values, respectively. The x- and y-axis represent the retention time and the m/z and in the background the total ion chromatogram (TIC) of the overlaid samples is shown.

**Fig. S2.**
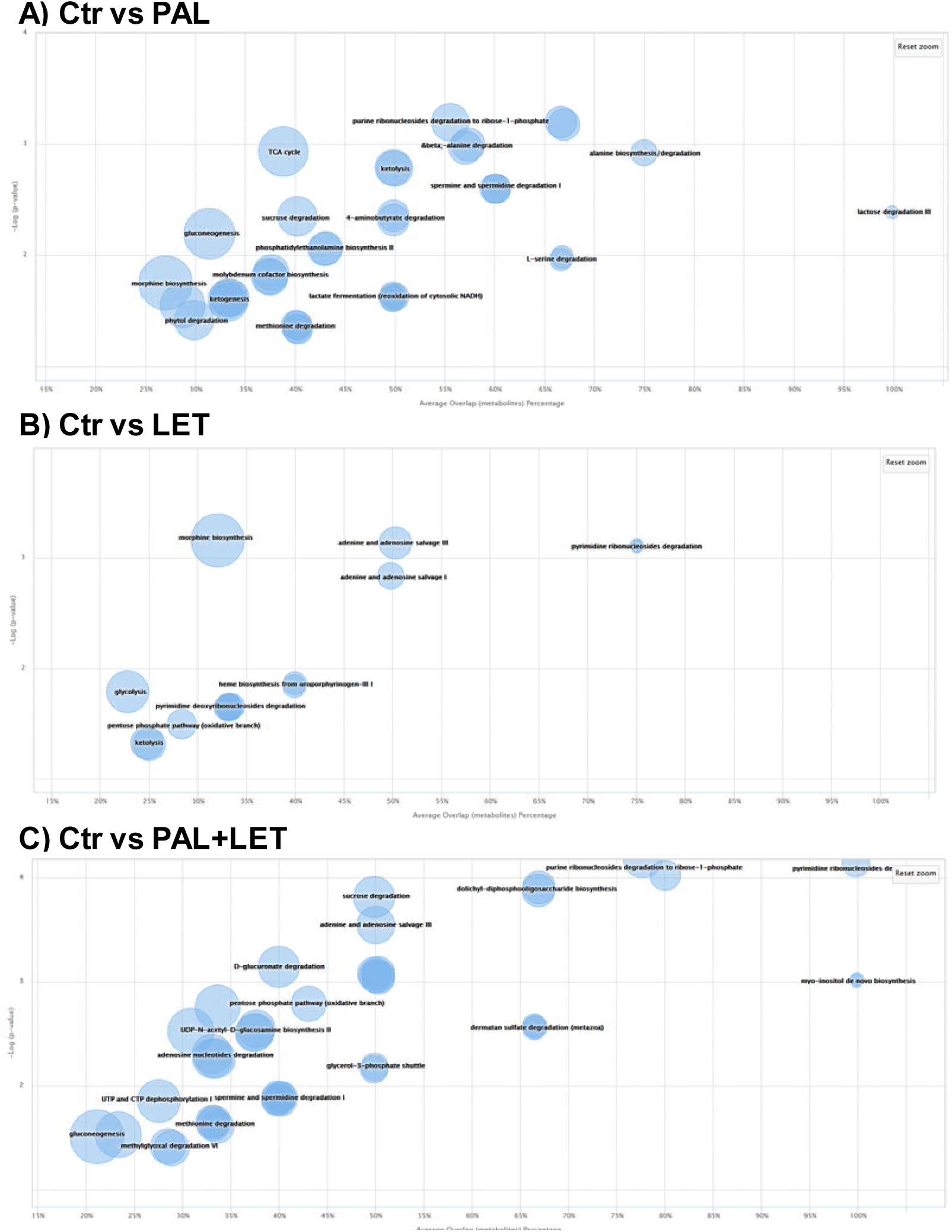
Predicted metabolic pathways highlighting the effect of the individual and combined agent dosing. Samples were generated after 48 h treatment with palbociclib (A), letrozole (B), and the combined treatment (C). Plots were generated using XCMS Online with the p-value threshold filter set to 0.05. The radius of a circle represents the size of the metabolic pathway; significantly dysregulated pathways appear in the upper right-hand quadrant of the plot. Pathways are plotted as a function of pathway significance versus average metabolic pathway overlap. The x- and y-axis represent the average metabolite overlap as percentage and the -log (p-value), respectively.

**Fig. S3.**
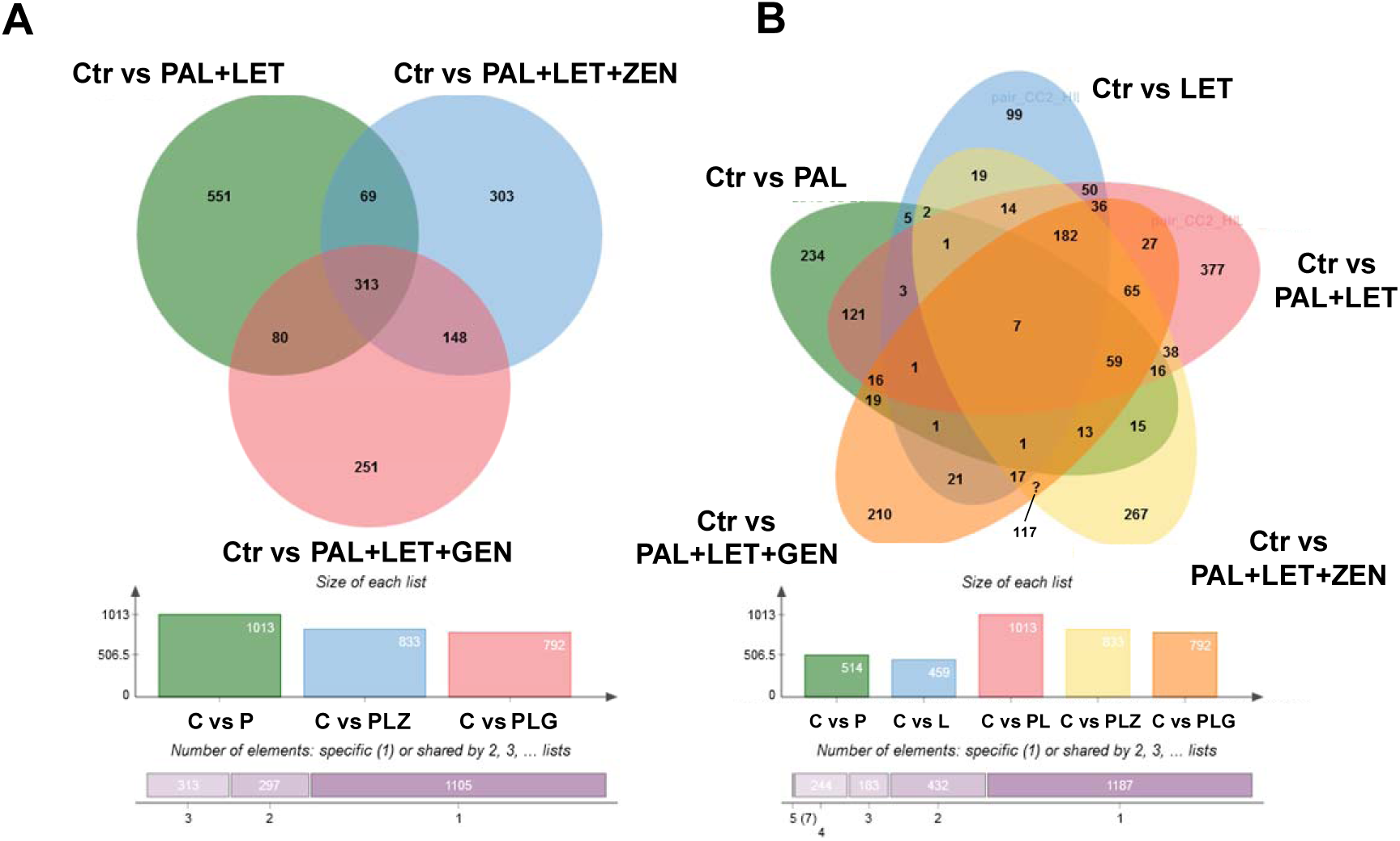
Venn diagrams of meta-analyses demonstrating the effect of xenoestrogens on the combined agents based on the HILIC-LC-MS measurements from cell extracts. (A) Compares the combined palbociclib and letrozole treatment alone (green) to cells exposed to ZEN (blue) or GEN (red) additionally. In (B) also the single agents were included in the analysis for the sake of completeness. Numbers indicate significantly dysregulated metabolic features compared to the control group (fold change >1.5, p-value <0.05).

## References

Adlercreutz, H. (2002). Phytoestrogens and breast cancer. Journal of Steroid Biochemistry and Molecular Biology 83, 113–118.

Agathocleous, M., and Harris, W.A. (2013). Metabolism in physiological cell proliferation and differentiation. Trends in Cell Biology 23, 484–492.

Beger, R.D., and Flynn, T.J. (2016). Pharmacometabolomics in drug safety and drug-exposome interactions. Metabolomics 12, 1–11.

Blei, T., Soukup, S.T., Schmalbach, K., Pudenz, M., Möller, F.J., Egert, B., Wörtz, N., Kurrat, A., Müller, D., Vollmer, G., et al. (2015). Dose-dependent effects of isoflavone exposure during early lifetime on the rat mammary gland: Studies on estrogen sensitivity, isoflavone metabolism, and DNA methylation. Molecular Nutrition and Food Research 59, 270–283.

Brasili, E., and Filho, V.C. (2017). Metabolomics of cancer cell cultures to assess the effects of dietary phytochemicals. Critical Reviews in Food Science and Nutrition 57, 1328–1339.

Cristofanilli, M., Turner, N.C., Bondarenko, I., Ro, J., Im, S.-A., Masuda, N., Colleoni, M., DeMichele, A., Loi, S., and Verma, S. (2016). Fulvestrant plus palbociclib versus fulvestrant plus placebo for treatment of hormone-receptor-positive, HER2-negative metastatic breast cancer that progressed on previous endocrine therapy (PALOMA-3): final analysis of the multicentre, double-blind, phase 3 randomised controlled trial. The Lancet Oncology 17, 425–439.

Dees, C., Foster, J.S., Ahamed, S., and Wimalasena, J. (1997). Dietary estrogens stimulate human breast cells to enter the cell cycle. Environmental Health Perspectives 105, 633–636.

Dhillon, S. (2015). Palbociclib: First Global Approval. Drugs 75, 543–551.

Finn, R., Dering, J., Conklin, D., Kalous, O., Cohen, D., Desai, A., Ginther, C., Atefi, M., Chen, I., Fowst, C., et al. (2009). PD 0332991, a selective cyclin D kinase 4/6 inhibitor, preferentially inhibits proliferation of luminal estrogen receptor-positive human breast cancer cell lines in vitro. Breast Cancer Research 11, R77.

Finn, R.S., Crown, J.P., Lang, I., Boer, K., Bondarenko, I.M., Kulyk, S.O., Ettl, J., Patel, R., Pinter, T., Schmidt, M., et al. (2015). The cyclin-dependent kinase 4/6 inhibitor palbociclib in combination with letrozole versus letrozole alone as first-line treatment of oestrogen receptor-positive, HER2-negative, advanced breast cancer (PALOMA-1/TRIO-18): a randomised phase 2 study. The Lancet Oncology 16, 25–35.

Finn, R.S., Martin, M., Rugo, H.S., Jones, S., Im, S.-A., Gelmon, K., Harbeck, N., Lipatov, O.N., Walshe, J.M., Moulder, S., et al. (2016). Palbociclib and Letrozole in Advanced Breast Cancer. New England Journal of Medicine 375, 1925–1936.

Forsberg, E., Huan, T., Rinehart, D., Benton, H.P., Warth, B., Hilmers, B., and Siuzdak, G. (2017). Data Processing, Pathway Mapping and Multi-Omic Systems Analysis using XCMS Online. Nature Protocols in press.

Goodson, W.H., 3rd, Luciani, M.G., Sayeed, S.A., Jaffee, I.M., Moore, D.H., 2nd, and Dairkee, S.H. (2011). Activation of the mTOR pathway by low levels of xenoestrogens in breast epithelial cells from high-risk women. Carcinogenesis 32, 1724–1733.

Hilakivi-Clarke, L., Onojafe, I., Raygada, M., Cho, E., Skaar, T., Russo, I., and Clarke, R. (1999). Prepubertal exposure to zearalenone or genistein reduces mammary tumorigenesis. Br J Cancer 80, 1682–1688.

Holliday, D.L., and Speirs, V. (2011). Choosing the right cell line for breast cancer research. Breast Cancer Research 13, 215.

Howlader, N., Altekruse, S.F., Li, C.I., Chen, V.W., Clarke, C.A., Ries, L.A.G., and Cronin, K.A. (2014). US Incidence of Breast Cancer Subtypes Defined by Joint Hormone Receptor and HER2 Status. JNCI: Journal of the National Cancer Institute 106, dju055–dju055.

Huan, T., Forsberg, E.M., Rinehart, D., Johnson, C.H., Ivanisevic, J., Benton, H.P., Fang, M., Aisporna, A., Hilmers, B., Poole, F.L., et al. (2017). Systems biology guided by XCMS Online metabolomics. Nat Meth 14, 461–462.

IARC (2014). World Cancer Report 2014, B W Stewart and C P Wild (The International Agency for Research on Cancer, WHO).

Ismail, A.M., In, L.L., Tasyriq, M., Syamsir, D.R., Awang, K., Mustafa, A.H., Idris, O.F., Fadl-Elmula, I., and Hasima, N. (2013). Extra virgin olive oil potentiates the effects of aromatase inhibitors via glutathione depletion in estrogen receptor-positive human breast cancer (MCF-7) cells. Food and chemical toxicology: an international journal published for the British Industrial Biological Research Association 62, 817–824.

Jäger, W., Gruber, A., Giessrigl, B., Krupitza, G., Szekeres, T., and Sonntag, D. (2011). Metabolomic Analysis of Resveratrol-Induced Effects in the Human Breast Cancer Cell Lines MCF-7 and MDA-MB-231. OMICS: A Journal of Integrative Biology 15, 9–14.

Jemal, A., Ward, E.M., Johnson, C.J., Cronin, K.A., Ma, J., Ryerson, A.B., Mariotto, A., Lake, A.J., Wilson, R., Sherman, R.L., et al. (2017). Annual Report to the Nation on the Status of Cancer, 1975– 2014, Featuring Survival. JNCI: Journal of the National Cancer Institute 109, djx030–djx030.

Johnson, C.H., Ivanisevic, J., Benton, H.P., and Siuzdak, G. (2015). Bioinformatics: The next frontier of metabolomics. Analytical Chemistry 87, 147–156.

Johnson, C.H., Ivanisevic, J., and Siuzdak, G. (2016a). Metabolomics: beyond biomarkers and towards mechanisms. Nat Rev Mol Cell Biol 17, 451–459.

Johnson, C.H., Patterson, A.D., Idle, J.R., and Gonzalez, F.J. (2012). Xenobiotic Metabolomics: Major Impact on the Metabolome. Annual Review of Pharmacology and Toxicology 52, 37–56.

Johnson, J., Thijssen, B., McDermott, U., Garnett, M., Wessels, L.F., and Bernards, R. (2016b). Targeting the RB-E2F pathway in breast cancer. Oncogene.

Jones, D.P. (2016). Sequencing the exposome: A call to action. Toxicology Reports 3, 29–45.

Ju, Y.H., Doerge, D.R., Woodling, K.A., Hartman, J.A., Kwak, J., and Helferich, W.G. (2008). Dietary genistein negates the inhibitory effect of letrozole on the growth of aromatase-expressing estrogen-dependent human breast cancer cells (MCF-7Ca) in vivo. Carcinogenesis 29, 2162–2168.

Lane, A.N., and Fan, T.W.M. (2015). Regulation of mammalian nucleotide metabolism and biosynthesis. Nucleic Acids Research 43, 2466–2485.

Mechcatie, E. (2015). FDA approves palbociclib with letrozole for advanced postmenopausal breast cancer. Oncology Report 11, 7.

Nagaraj, G., and Ma, C. (2015). Revisiting the estrogen receptor pathway and its role in endocrine therapy for postmenopausal women with estrogen receptor-positive metastatic breast cancer. Breast Cancer Res Treat 150, 231–242.

NCI (2017). National Cancer Institute SEER Website Cancer Statistics (https://seer.cancer.gov/faststats/selections.php, April 30th 2017).

Nechuta, S.J., Caan, B.J., Chen, W.Y., Lu, W., Chen, Z., Kwan, M.L., Flatt, S.W., Zheng, Y., Zheng, W., Pierce, J.P., et al. (2012). Soy food intake after diagnosis of breast cancer and survival: an in-depth analysis of combined evidence from cohort studies of US and Chinese women. The American Journal of Clinical Nutrition 96, 123–132.

Patti, G.J., Tautenhahn, R., and Siuzdak, G. (2012a). Meta-analysis of untargeted metabolomic data from multiple profiling experiments. Nat Protocols 7, 508–516.

Patti, G.J., Yanes, O., and Siuzdak, G. (2012b). Innovation: Metabolomics: the apogee of the omics trilogy. Nat Rev Mol Cell Biol 13, 263–269.

Pazaiti, A., Kontos, M., and Fentiman, I.S. (2012). ZEN and the art of breast health maintenance. International journal of clinical practice 66, 28–36.

Rawat, A., Gopal, G., Selvaluxmy, G., and Rajkumar, T. (2013). Inhibition of ubiquitin conjugating enzyme UBE2C reduces proliferation and sensitizes breast cancer cells to radiation, doxorubicin, tamoxifen and letrozole. Cellular Oncology 36, 459–467.

Rice, S., and Whitehead, S.A. (2006). Phytoestrogens and breast cancer –promoters or protectors? Endocrine-Related Cancer 13, 995–1015.

Saxton, R.A., and Sabatini, D.M. (2017). mTOR Signaling in Growth, Metabolism, and Disease. Cell 168, 960–976.

Shanle, E.K., and Xu, W. (2011). Endocrine disrupting chemicals targeting estrogen receptor signaling: identification and mechanisms of action. Chem Res Toxicol 24, 6–19.

Smith, C.A., O'Maille, G., Want, E.J., Qin, C., Trauger, S.A., Brandon, T.R., Custodio, D.E., Abagyan, R., and Siuzdak, G. (2005). METLIN: a metabolite mass spectral database. Therapeutic drug monitoring 27, 747–751.

Soukup, S.T., Helppi, J., Müller, D.R., Zierau, O., Watzl, B., Vollmer, G., Diel, P., Bub, A., and Kulling, S.E. (2016). Phase II metabolism of the soy isoflavones genistein and daidzein in humans, rats and mice: a cross-species and sex comparison. Archives of Toxicology 90, 1335–1347.

Sullivan, M.R., and Vander Heiden, M.G. (2017). When cancer needs what's non-essential. Nat Cell Biol 19, 418–420.

Takemura, H., Shim, J.-Y., Sayama, K., Tsubura, A., Zhu, B.T., and Shimoi, K. (2007). Characterization of the estrogenic activities of zearalenone and zeranol in vivo and in vitro. The Journal of Steroid Biochemistry and Molecular Biology 103, 170–177.

Tautenhahn, R., Patti, G.J., Rinehart, D., and Siuzdak, G. (2012). XCMS Online: A Web-Based Platform to Process Untargeted Metabolomic Data. Analytical Chemistry 84, 5035–5039.

Vejdovszky, K., Schmidt, V., Warth, B., and Marko, D. (2017). Combinatory estrogenic effects between the isoflavone genistein and the mycotoxins zearalenone and alternariol in vitro. Molecular Nutrition & Food Research 61, 1600526.

Verkasalo, P.K., Appleby, P.N., Allen, N.E., Davey, G., Adlercreutz, H., and Key, T.J. (2007). Soya intake and plasma concentrations of daidzein and genistein: validity of dietary assessment among eighty British women (Oxford arm of the European Prospective Investigation into Cancer and Nutrition). British Journal of Nutrition 86, 415–421.

Warth, B., Levin, N., Rinehart, D., Teijaro, J., Benton, H.P., and Siuzdak, G. (2017a). Metabolizing Data in the Cloud. Trends in Biotechnology 35, 481–483.

Warth, B., Spangler, S., Fang, M., Johnson, C., Forsberg, E., Granados, A., Martin, R.L., Domingo, X., Huan, T., Rinehart, D., et al. (2017b). Exposome-Scale Investigations Guided by Global Metabolomics, Pathway Analysis, and Cognitive Computing. Analytical Chemistry in press.

Warth, B., Sulyok, M., Berthiller, F., Schuhmacher, R., and Krska, R. (2013). New insights into the human metabolism of the Fusarium mycotoxins deoxynivalenol and zearalenone. Toxicology Letters 220, 88–94.

Wolff, A.C. (2016). CDK4 and CDK6 Inhibition in Breast Cancer — A New Standard. New England Journal of Medicine 375, 1993–1994.

Yager, J.D., and Davidson, N.E. (2006). Estrogen Carcinogenesis in Breast Cancer. New England Journal of Medicine 354, 270–282.

Zamboni, N., Saghatelian, A., and Patti, Gary J. (2015). Defining the Metabolome: Size, Flux, and Regulation. Molecular Cell 58, 699–706.

Zhang, F.F., Haslam, D.E., Terry, M.B., Knight, J.A., Andrulis, I.L., Daly, M.B., Buys, S.S., and John, E.M. (2017). Dietary isoflavone intake and all-cause mortality in breast cancer survivors: The Breast Cancer Family Registry. Cancer 123, 2070–2079.

